# Patterns of Ocular Dominance are in the Eye of the Beholder

**DOI:** 10.64898/2025.12.14.694226

**Authors:** Spencer Rooke, Seoyeon Choi, Vijay Balasubramanian

## Abstract

Ocular Dominance Columns (ODCs), structures in the early visual cortex that demarcate inputs from each eye, have been observed in a wide range of mammals. The extent and arrangement of these columnar structures can vary drastically from species to species, and even along the visual cortex of a single individual. While previous studies show that these structures can form by competitive Hebbian learning, the relationship between system parameters and the resulting organizational patterns remains unclear. Here, we use a mesoscopic model of V1 development to explain how the different observed patterns of ocular dominance (columns, islands, and monocular) can arise through local Hebbian competition, as a function of the density of projections from each eye and cortical interaction strengths. We argue that our results explain the broad cortical organizational differences between species which have forward-facing versus steeply angled eyes. We additionally show that spatially varying retinal input projections, which can arise from a combination of retinotopy and varying cell density, can lead to coexistence of multiple ocular dominance patterns along the cortical sheet, consistent with experimental observations.

## I. INTRODUCTION

In mammals, visual inputs are received by photoreceptors in the retina of each eye, where they are processed and compressed by retinal ganglion cells (RGCs) into neural signals. These signals are sent down optic nerves associated with each eye, passing through the lateral geniculate nucleus (LGN), and thence to the primary visual cortex (V1) located in the occipital lobe ([1], Fig.1). Each half of the mammalian visual cortex typically responds to projections from opposing visual fields; that is, stimuli in the left visual field will elicit responses in the right visual cortex, and likewise, stimuli in the right visual field will elicit responses in the left visual cortex [2]. Both eyes receive visual input from both the left and right visual field; thus both halves of V1 receive projections from both eyes. As a result, each half of the primary visual cortex, one of the earliest stages of signal processing in the brain, must consistently integrate inputs from each eye.

**FIG. 1.**
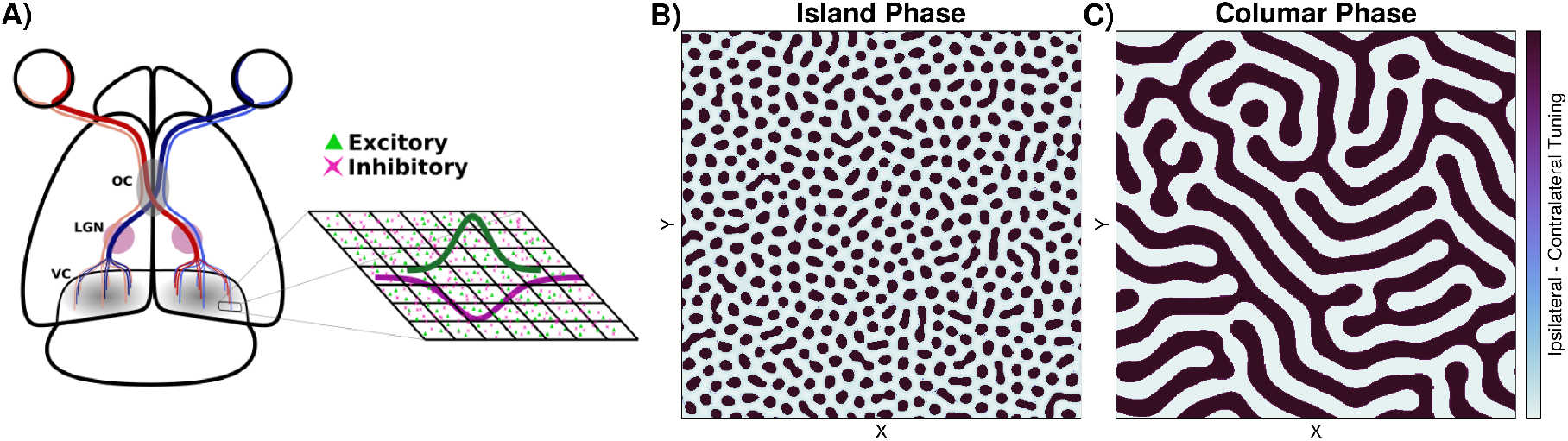
**(A)** Visual pathway of the rodent, with the ocular chasm (OC), lateral geniculate nucleus (LGN), and visual cortex (VC) marked. We adopt a mesoscopic view of activity along V1, where the total neural activity *r* about a position of *X* can be viewed as a local average over the activity of neurons on the neural sheet. Assuming local interactions (i.e., at least exponential falloff with distance), and that inhibition has a longer associated length scale than excitation, yields an interaction kernel that is a difference of gaussians. **(B-C)** Demonstration of two of the ordered phases: islands, and the columnar phase. Not shown are the monocular phase and the disordered phases.

As first observed by Hubel and Wiesel in cats [3], cells along the cortical sheet of V1 will respond preferentially to one eye or another. This preferential tuning is in many mammals not randomly distributed across the cortical sheet. Rather, nearby cells along the cortical sheet often have similar tunings. It was later observed that in macaques, this preferential response occurs in bands along the surface of portions of the striate cortex [4–6]. These bands, known as Ocular Dominance Columns (ODCs), have since been observed in a variety of mammals including macaques, cats, ferrets, and humans [3, 5–9]. While less pronounced and not appearing in bands, related structures have additionally been observed in rodents [10]. The extent and arrangement of these columnar structures can vary drastically from species to species, and even along the visual cortex of a single animal [3, 5–9]. Notably, mammals with prominent columns in their central field of view are typically predators, with two forward-facing eyes and large visual overlap, while those with less pronounced or absent columns typically have steeply angled eyes [11]. Thus, some have speculated that the formation of ocular dominance columns depends on the relative angle of the eyes, and is necessary for stereopsis [11–13], as this would result in a large asymmetry in the visual input received from each eye along the cortical sheet. We will explore this hypothesis here.

To understand the formation of ODCs, we consider a mesoscopic model of neural activity along the cortical sheet of the primary visual cortex (V1), in which neurons locally synapse onto each other, while additionally receiving inputs from the LGN, which in turn originate from upstream visual input. We will suppose that recurrent connections remain fixed, while the feedforward connections from upstream visual areas are dynamic and update via Hebbian learning. Previous theoretical work has demonstrated that Hebbian plasticity can give rise to both ODCs and orientation selectivity along the cortical sheet [14–19].

We build on previous work by incorporating the relative density of upstream retinal input from each eye synapsing onto V1. This leads to find three distinct regimes of self-organized ocular dominance – columns, islands, and monocular – that depend on the relative contribution of each eye, and tuned by the cortical interaction strength. ODC formation is known to occur before eye opening and during early stages of development [12]. We use a simplified model of random retinal dynamics, with local instantaneous spatial correlations that mimic those found in normal mice [20].

We additionally incorporate spatially varying input densities, to mimic variations of inputs along the cortex that can arise from retinotopy and differences in retinal ganglion cell densities. By constructing these densities in a manner consistent with the relative number of retinal photoreceptive cells and the retinotopy induced by the relative eye opening angle, we observe coexistence of multiple ocular dominance patterns on a single cortical sheet. In particular, regions with high visual overlap exhibit varying degrees of binocular coexistence, which shrink and vanish into the periphery. This qualitatively matches what is seen in many mammals. For example, in cats, macaques, and humans, distinct columns form over a large portion of the visual cortex in regions of high visual overlap, but shrink into islands then vanish in the periphery [3, 5–9], while rodents have small islands of binocular coexistence, and large regions of monocular dominance along their visual cortex [10]. This lends support to the idea that the relative density of afferent fibres, which is partially set by eye-opening angle through retinotopy, is an important driving force in the development of the observed patterns of ocular dominance in V1.

### A. Model Description

We consider a simple mesoscopic model of neural activity along each half of the visual cortex (V1), with dynamics:

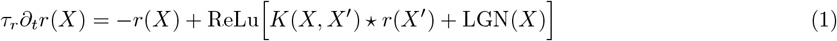

Here, ReLu denotes a rectified linear transformation (namely negative arguments are set to zero), “⋆” is the convolution operation, *X* denotes position along the cortical sheet, *r*(*X*) is the average firing rate of neurons about that position, and τ is the timescale of response to an input pulse. That is, *r*(*X*) does not represent the firing rates of a single neuron, but rather represents the coarse grained activity of populations of neurons located about *X*. Neurons in the visual cortex receive feedforward input from the LGN and recurrent input from other neurons along the cortical sheet, modeled here by a recurrent interaction kernel *K*. A rectifying nonlinearity enforces positive activity. The intracortical interaction kernel is given by a difference of Gaussians (DoG) (Figure 1)

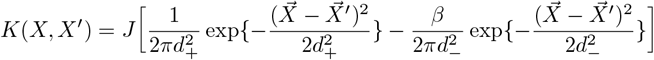

Here, *d*_+_ and *d*_−_ set the excitatory and inhibitory interaction distance, *β* sets the relative influence of inhibitory and excitatory activity, and *J* sets the relative importance of recurrent interactions on the cortical sheet compared to feedforward activity from each eye. The DoG kernel can be justified in the following way. Typical neural interactions along the cortical sheet are local, and so the interaction kernel can be taken to fall off exponentially with some function of distance along the cortical sheet. The squared distance is a natural choice and often assumed [14, 15, 19, 21]. Additionally a common neural circuit motif is that excitatory neurons synapse more locally than inhibitory neurons, but with greater efficacy, so that the length scale associated with the inhibitory interaction should be larger than that associated with the excitatory interaction, *d*_−_ > *d*_+_, leading to the difference of gaussian kernel given above. This form is often assumed theoretically [14, 15, 19, 21], and while the exact form of recurrent connectivity is not known, experimental evidence support the approximate description used here [22–24]. Other forms for the connectivity with short range excitation and long range inhibition lead to phenomenologically similar behaviour [14]. Overall, the dynamics is effectively a coarse grained version of a standard recurrent rate model *τ*∂_*t*_*r*_*i*_ = −*r*_*i*_ + ReLu[*J*_*ij*_*r*_*j*_ + *h*_*i*_], coarse-grained in space, with some assumptions about the structure of cortical connectivity.

Neurons additionally receive feedforward input from the LGN. In most mammals, the LGN does not mix the signals coming from each eye [25]. As such, one can split the neural signals coming from the LGN into ipsilateral and contralateral components, based on retinal signal origin, so that LGN(*X*) = *W*_*I*_ (*X*)*ρ*_*I*_ (*X*)*I*_*I*_ (*X*)+*W*_*C*_(*X*)*ρ*_*C*_(*X*)*I*_*C*_(*X*). Here *I*_*I/C*_ reflects retinal activity of the Ipsilateral or Contralateral eye, *W*_*I/C*_ reflects the relative synaptic strength of LGN projections onto *V* 1, and *ρ*_*I/C*_ acts as a transfer function capturing effects of varying retinal signal strength arising from a combination of retinotopy and varying retinal fibre density. The neural activity along the cortical sheet can thus be represented by the differential equation:

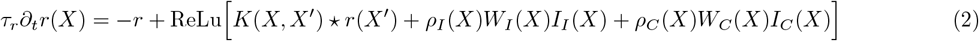

Note that *W*_*I*_ (*X*) and *W*_*C*_(*X*) specify how the neural response *r*(*X*) is tuned; if *W*_*I*_ is large then r will predominantly respond to inputs from the ipsilateral eye, and likewise, if *W*_*C*_ is large then r will respond to inputs from the contralateral eye. During development, these connectivities are plastic, and we model their plasticity with a simple Hebbian rule:

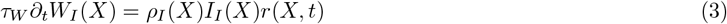

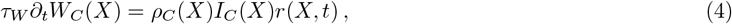

where *τ*_*W*_ represents the learning rate, *τ*_*W*_ >> *τ*_*r*_, and *r*(*X, t*) depends implicitly on the weights, and obeys Eq. (2). We additionally assume the total number of afferents onto *V*1 is bounded at each position X, so that 0 ≤ *W*_*I*_ (*X*) + *W*_*C*_(*X*) ≤ 1, which reflects limited synaptic real estate around *X*. Competition is crucial for the proper development of ODCs via Hebbian plasticity, and this bound on the total number of afferents introduces competition in a natural way. In particular, if *W*_*I*_ (*X*) + *W*_*C*_(*X*) = 1 saturates this bound for the duration of the learning, then the learning dynamics can be written in the explicitly competitive form:

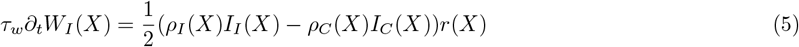

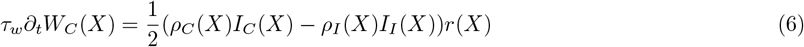

This model simplifies the more detailed models in [14–17, 26] while keeping the essential features, to enable exploration of the parameter space and analysis of the effects on pattern formation. At intermediate timescales *τ*_*r*_ < *t* < *τ*_*I*_, the dynamics reduce to the self consistency conditions described by the one dimensional model found in [21].

## II. RESULTS

Having established a model of cortical activity and development, we explored the effects that parameter variation has on the system. Before eye opening, we model spontaneous retinal activity, which is known to spur columnar development, as independent between the two eyes and evolving randomly in time. For simplicity, we assume a minimal model of random activity for the retinal neurons/input, given by the Stochastic Differential Equations:

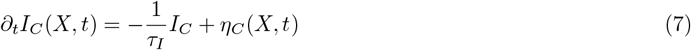

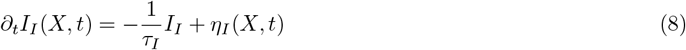

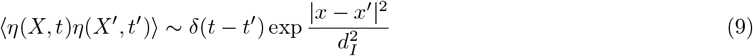

Here, *η*(*X, t*) is temporally white but spatially colored noise with length scale *d*_*I*_, consistent with known instantaneous statistics for the (wild-type) mouse visual system [20], and *τ*_*I*_ sets the time scale of retinal activity evolution. For simplicity, we do not model the detailed dynamics of retinal waves, but rather use minimal stochastic dynamics that mimic known low order statistics. A single instance of the mapped retinal activity (before modulation by *ρ*) is then a time evolving gaussian random field with the following correlation structure:

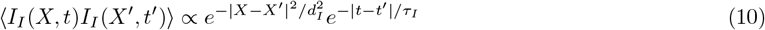

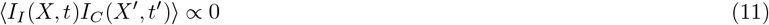

From numerical experiments, we find that structures along the cortical sheet only form when *τ*_*I*_ << *τ*_*W*_ and *d*_*I*_ > *d*_−_ > *d*_+_. Intuitively, the time scale of the learning must be slower than the rate at which inputs change or learning fail to converge. The length scale of input correlations also must be sufficiently large for structures to form, a point which we will return to in what follows. It is also worth noting that increasing the overall input strength has the same effect as increasing the learning rate *τ*_*W*_ and decreasing the cortical interaction strength *J*, and so for simplicity we set proportionality constants to one. I The remaining parameters that affect the system include the cortical length scales *d*_*±*_, the cortical interaction strength *J*, and the relative densities *ρ*_*I/C*_. To explore the phase structure associated with these densities, we first consider the case where they are not spatially varying. By simulating the dynamics of Eq. (6) one finds that the ratio *η* ≡ *ρ*_*I*_ /*ρ*_*C*_ and the cortical strength *J* are primarily responsible for setting the phase structure. We observed three ordered phases under variations of these parameters, which correspond to a “columnar phase” with roughly equal ipsilateral and contralateral representation, an “island” phase with small ipsilateral islands in a contralateral background, and a phase associated with total monocular dominance. The first two phases are shown in Fig. 1,2.

**FIG. 2.**
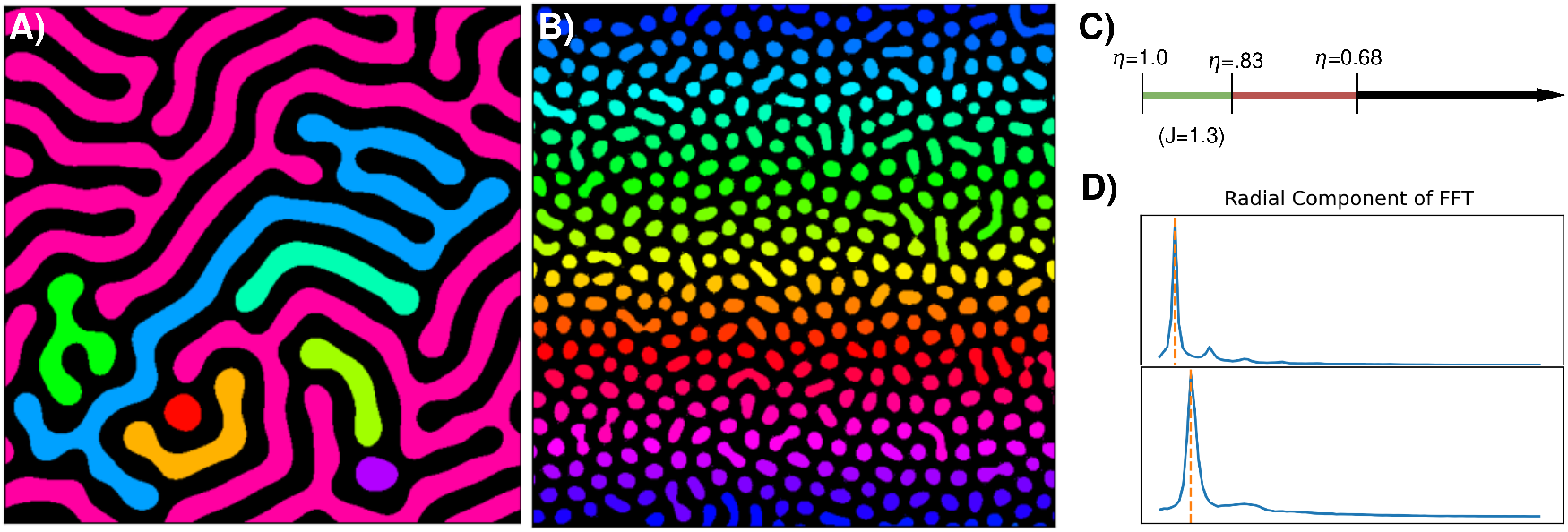
**(A-B)** The two nontrivial phases, as previously described. Colors indicate clusters identified using invasion percolation [27, 28]. **(C)** As the ratio the relative contralateral contribution to activity on the neural sheet shrinks (*ρ*_*C*_*/ρ*_*I*_ shrinks) the system exhibits a phase transition from the columnar phase to the island phase, and then to monocular dominance. **(D)** The radial component of the fourier transform in the **(Top)** columnar phase and **(Bottom)** in the island phase. The peak peaks set the characteristic length scale of observed structure.

Conversely, the length scales *d*_*±*_ set the maximum characteristic width of observed structures. To see why, we consider Eq. (2) over a region Ω_+_ where activity is positive, so that the dynamics is linear. For time scales *t* >> *τ*_*r*_ and *t* << *τ*_*W*_ the dynamics equilibrate, and one has that on this region:

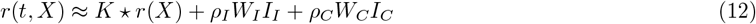

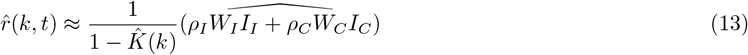

Here we used the Fourier transform to turn the convolution in (12) into a product; thus the hatted quantities in Eq. 13 such 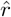 are Fourier transforms of the corresponding quantities in (12). For simplicity consider densities *ρ*_*I/C*_ and inputs *I*_*I/C*_ which are not spatially varying. Writing Eq. (6) in fourier space and inserting the above equation for the rates yields:

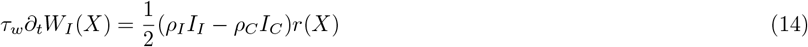

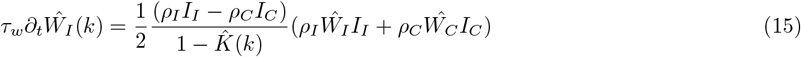

Structures of characteristic size *λ* = 2*π*/|*k*| are thus promoted if *ρ*_*I*_ *I*_*I*_ − *ρ*_*C*_*I*_*C*_ > 1. Furthermore, the fastest growing structures will typically be those whose wavenumber maximizes 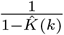. These are wavenumbers with magnitude:

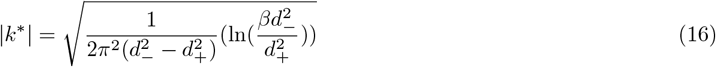

As such, the most promoted structures have a characteristic width ∼ π/ |*k*^∗^|. Further, the DC component associated with monocular dominance can be suppressed by taking *β* > 1. This suggests that in the balanced / columnar phase, the local spatial scale is at least partially determined by the anatomy of the visual cortex, and less dependent on visual input, which was verified in simulation of Eq. (6). However, the 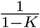 term is flat for frequencies larger than |*k*^∗^|, and so peaks in the Fourier spectrum in the inputs for |*k* |> |*k*^∗^| that persist in time could potentially disrupt this natural length scale set by the system. Further, we have assumed for simplicity that there are large domains Ω_+_ of activity at any given time, but if typical domain sizes are smaller than this, then structures with this natural characteristic length scale may fail to dominate. This may partially explain why the spatial scale of the random retinal input *d*_*I*_ must be sufficiently large for structures to form.

We now turn to exploring the phase structure associated with variation of the cortical interaction strength *J* and the relative input ratios *η* = *ρ*_*I*_ /*ρ*_*C*_. For a given set of parameters, we simulate multiple trajectories of Eq. (6). Each trajectory produces a set of learned synaptic weights *W*_*I/C*_ (*X*) along the cortical sheet, from which we can extract measures associated with each phase. When structures form, the bound *W*_*I*_ + *W*_*C*_ ≤ 1 is typically fully saturated by *W*_*I*_ or *W*_*C*_, and so clusters of ipsilateral and contralateral afferents are well defined.

To characterize the spatial organization, we use invasion percolation to compute the largest cluster size, the number of distinct clusters, and the total Ipsilateral-Contralateral alignment. As shown in Figure 2, the number of identified clusters is small in the columnar phase, large in the island phase, and vanishing in the monocular phase. Similarly, the largest cluster size shrinks in the island phase and vanishes in the monocular phase, while the total Ipsilateral-Contralateral allignment grows in the island phase and saturates in the monocular phase. As such, these three measures act as good order parameters for identifying the observed phase.

As these order parameters vary smoothly across the transitions, we identify peaks in their second derivative averaged over many learning trajectories to identify phase boundaries. This lets us characterize the organization of the cortical network as a function of input balance, with phase boundaries drawn in Figure 2. In regions with balanced inputs *ρ*_*I*_ ∼*ρ*_*C*_, we typically observe well defined columns. Reducing this balance so that *ρ*_*I*_ < *ρ*_*C*_ causes the columns to shrink into the island phase, with further reduction causing the disappearance of ipsilateral islands and the onset of the monocular phase.

In the visual cortex of a mammal, we expect *ρ*_*I/C*_ to vary with position *X* over the cortical sheet. For example, in the left visual cortex, the representation of the right visual field depends on the number of signals arriving from each eye at each position *X*. As such, *ρ*_*I/C*_ effectively act as transfer functions for signals originating in the visual field, and is shaped by an animal’s anatomy. This means that factors such as retinotopy, the density of retinal ganglion cells with respect to eye eccentricity, and eye opening angle are expected to effect *ρ*_*I/C*_ directly, as shown in Figure 3.

**FIG. 3.**
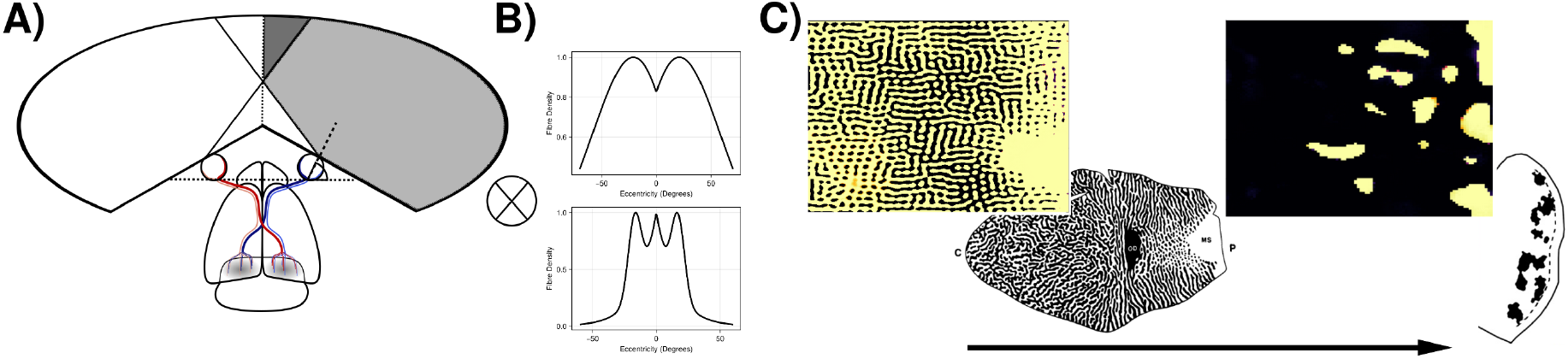
**(A)** The rodent visual system, with the right visual field identified. The eye opening angle determines the degree of visual overlap. Combined with **(B)** retinal ganglion cell densities, one can construct an approximation of the densities *ρ*_*I*_*/C*(*X*) for the left visual cortex. **(C) (Left)** With shallow eye opening angle, one finds large regions of the columnar phase, with the island and monocular phase appearing in the periphery, phenomenologically recreating the phase coexistence observed in the human visual cortex. **(Right)** With steep eye opening angle, monocular tuning dominates due to the lack of visual overlap. We observe islands in the regions associated with the small region of visual overlap, phenomenologically recreating the phase coexistence observed in the rodent visual cortex. Stains recreated from [9, 10] with permission.

In animals with two forward facing eyes, one expects large regions of the visual cortex with *ρ*_*I*_ ∼ *ρ*_*C*_, corresponding to regions with large visual overlap, and smaller peripheral regions where visual overlap shrinks and vanishes. In these regions, we likewise expect the columnar phase to fall off, and give way to an island or monocular phase. This is typical, for example, in cats, humans, and macaques [3, 5–7, 9], and this pattern is reproduced in simulations of Eq. (6), as shown in Figure 3, where gradients in *ρ*_*I*_ and *ρ*_*C*_ naturally reproduce the phase coexistence observed across a single cortical sheet. Conversely, with eyes that are significantly angled outwards, visual overlap is limited, and contralateral inputs largely dominate. Across much of the cortical sheet, *ρ*_*C*_ will thus dominate *ρ*_*I*_. Simulating Eq. (6) in this scenario leads to large monocular regions, with only small regions of potential binocular coexistence appearing in the regions associated with visual overlap (Figure 3). This is consistent with measurements in the rodent visual cortex [10].

## III. DISCUSSION

In this work, we studied how cortical interactions and visual input densities can combine to shape the self-organization of ocular dominance patterns in the primary visual cortex. We suggest that various patterns of ocular dominance seen in mammals depend primarily on relative fiber densities from LGN associated with each eye, which in turn depend on a combination of retinotopy and retinal fiber densities. We demonstrate that variations in these parameters along with simple Hebbian dynamics can lead to the development of three experimentally observed structures: ocular dominance columns, islands, and monocular organization. This result relies on the the fact that the LGN maintains segregated inputs from each eye [25]. While there are exceptions in portions of the rodent LGN [29], this framework still provides a simple explanation for ocular dominance diversity. We additionally show that in the columnar phase, the characteristic width is primarily set by the intercortical interaction kernel, which acts as a weak filter promoting wavelengths dependent on the excitory and inhibitory interaction distances *d*_*±*_. This implies that the spatial scale of the observed structures, and in particular columns, is partially set by the anatomy of the cortex. This idea can be experimentally tested, and may be consistent with experimental evidence relating stripe patterns and cortical geometry [30, 31]. Finally, we explored the role of spatially varying retinal densities, and demonstrate that this can lead to coexistence of different patterns of ocular dominance along the cortical sheet which phenomenologically matches observations [3, 5–7, 9, 10].

While our simple framework captures key aspects of ocular dominance development, it omits some biological details. We consider only local, non-critical period retinal activity statistics, only model the overall correlations (and not the detailed dynamics) of retinal waves, and do not consider activity dynamics after eye opening. We also neglect orientation selectivity, which is thought to play a role in anchoring ODCs [26, 32–34]. Despite these omissions or minimal model broadly reproduced the observed phenomenology of ocular dominance in visual cortex. Extending our model to include more biologically realistic transfer functions, the effects of plasticity after eye opening, and learned orientation tuning may allow a more fine-grained, quantitative match with structures seen across different species.

## Funding

This work was supported in part by the NSF and DoD OUSD (R & E) under Agreement PHY-2229929 (The NSF AI Institute for Artificial and Natural Intelligence). VB was supported in part by the Eastman Professorship at Balliol College, Oxford.

## Notes

### Competing Interest Statement

The authors have declared no competing interest.

